# Probing the glioma micro-environment: analysis using biopsy in combination with ultra-fast cyclic immunolabeling

**DOI:** 10.1101/2024.06.15.599078

**Authors:** Thomas S. van Solinge, Juhyun Oh, Erik Abels, Peter Koch, Xandra O. Breakefield, Ralph Weissleder, Marike L.D. Broekman

## Abstract

The interaction between gliomas and the immune system is poorly understood and thus hindering development of effective immunotherapies for glioma patients. The immune response is highly variable during tumor development, and affected by therapies such as surgery, radiation, and chemotherapy. Currently, analysis of these local changes is difficult due to poor accessibility of the tumor and high-morbidity of sampling. In this study, we developed a model for repeat-biopsy in mice to study these local immunological changes over time. Using fine needle biopsy we were able to safely and repeatedly collect cells from intracranial tumors in mice. Ultra-fast cycling technology (FAST) was used for multi-cycle immunofluorescence of retrieved cells, and provided insights in the changing immune response over time. The combination of these techniques can be utilized to study changes in the immune response in glioma or other intracranial diseases over time, and in response to treatment within the same animal.

**Teaser:** Fine-needle biopsy and ultra-fast cycling technology techniques were developed to allow for repeat sampling and analysis of glial tumors in mice.

## Introduction

Glial tumors are malignant intra-axial brain tumors known for their aggressive growth and resistance to treatment, with abysmal survival rates^1,2^. To accurately diagnose and subtype gliomas, tissue is needed^3^. Forms of liquid biopsy such as blood or CSF-based tests can provide information about the status of tumors, but are thus far not able to consistently and accurately diagnose or monitor glioma in a clinical setting^4^. With the increasing interest in the interaction between tumor and its microenvironment, isolation of tumor and local immune cells has only grown in importance. In patients, tumor tissue is acquired during (re-) resection, or via stereotactic biopsy. These procedures carry risk, which increases with each subsequent surgical intervention^5^. Additionally, it is well-documented that glioblastomas are heterogeneous in nature, with a large variability in (epi)genetic, molecular, and phenotypic characteristics between glioma cells within the same tumor^6^. Not only is there heterogeneity spatially within the tumor^7^, the molecular traits of glioblastoma can change over time, especially in response to treatments, such as surgery, chemotherapy, and radiation^8^. With the tumor, the microenvironment and the immune response changing as well, this leads to different immune profiles and potential targets for immunotherapy over time^9^.

While we know glioblastoma itself and the immune response are changing over time, in patients we are currently only able to measure this at crude intervals: at the initial surgery, and at subsequent re-resection(s) upon treatment failure. There have been calls to perform stereotactic biopsies more often for personalized therapy and research purposes^10^, but this is rarely done in practice due to the burden on patients, and the increase in mortality and morbidity with each additional biopsy^5^.

Animal models provide more flexibility in analysis of molecular and immunological changes during disease progression. Mouse brain tumor models, while known for various short-comings^11^, have been used to demonstrate the impact of therapy on the tumor microenvironment^12,13^. However, current models require euthanization of the animal to study the molecular and immunological landscape in depth, and thus require many animals to evaluate changes over time. Furthermore, comparing different time points from different animals introduces a major source of variability, making it difficult to find subtle changes in the tumor or its microenvironment. Intravital imaging allows for the monitoring of cell movement *in vivo* over time, but this method only allows for a handful of cell types to be studied at the same time^14^.

In this study, we present a model to safely repeat biopsies in a mouse model of glioblastoma and accurately analyze the immune profile of the tumor microenvironment. Using a stereotactic frame and minimally invasive biopsy techniques, we were able to retrieve cells from the brain repeatedly, up to 3 times. To allow for in-depth analysis of the immune profile with a limited number of cells, we applied multiplexed cellular fluorescent imaging. Ultra-fast cycling technology (FAST) was utilized for multi-cycle immunofluorescence in single cells. We validated our results with flow cytometry, and we were able to show changes in the immune profile over time. Finally, we identified small but distinct changes in the local immune response after biopsy. This repeat biopsy assay has the potential to serve as a useful tool for studying the complex and rapidly evolving tumor microenvironment during cancer treatment.

## Results

### Repeated brain biopsy is safe and feasible in mice with brain tumors

For development of this technique, we implanted C57BL/6 mice with GL261.Fluc.GFP tumors^15^. This syngeneic tumor model is commonly used to monitor glioma *in vivo* and the immune response is well characterized^11^. Tumors were implanted in the striatum at day 0. At day 10, IVIS imaging was performed to confirm tumor growth, and mice were stratified into two groups of equal average tumor size. One group received a biopsy at the original implantation site with a 22s gauge Hamilton syringe on day 10, and another biopsy on day 15 after implantation (*Figure 1A)*. About 10-20 ul of tissue was removed, equating to between 7000 and 15.000 cells. The control group received a sham surgery at the same time-points, where the previous incision was opened and closed, but no biopsy performed. Both groups received the same duration of anesthesia and post-operative care. There was no difference in overall survival (OS) between these two groups (*Figure 1B),* showing that our technique was safe and that biopsy itself did not lead to slower or increased tumor growth.

**Figure 1:**
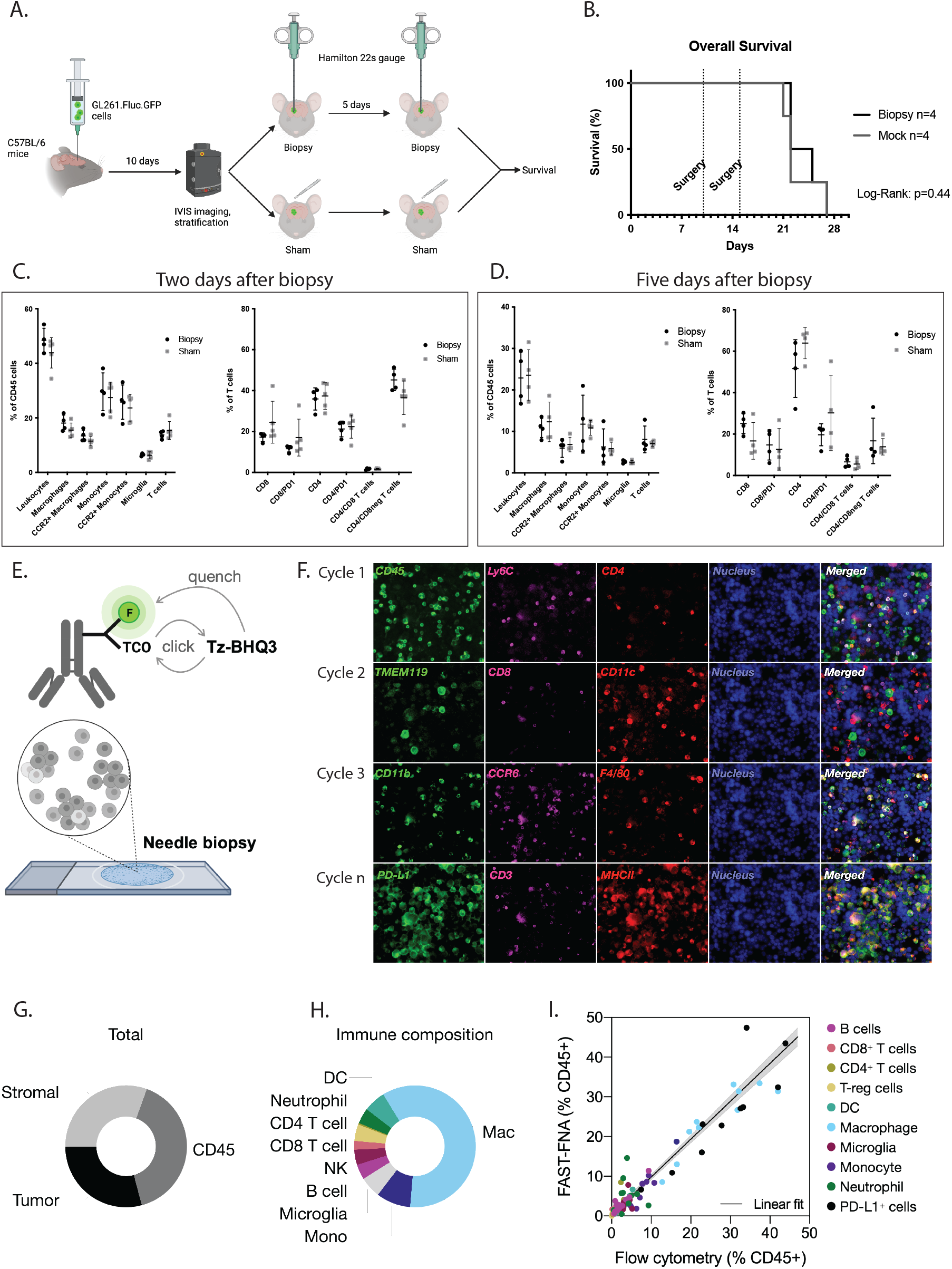
Serial sampling in mice is safe and accurate. **A)** Flow chart for survival experiment. One group received repeat biopsy, one group received sham-surgery. **B)** Kaplan-Meier curve comparing biopsied versus non-biopsied mice with brain tumors. Two groups of 4. Log-Rank test, p=0.44. **C)** Analysis of immune populations studied by flow cytometry on tumor containing hemispheres in biopsied versus non-biopsied mice after two days. **D)** Analysis of immune populations studied by flow cytometry on tumor containing hemispheres in biopsied versus non-biopsied mice after five days. **E)** schematic representation of the FAST methodology for staining. **F)** Example of repeat imaging using FAST technology on cells obtained through brain tumor biopsy. **F)** Example of phenotyping of cells obtained through brain tumor biopsy. **G)** Example of phenotyping of cells obtained through brain tumor biopsy. **H)** Example of phenotyping of immune cells obtained through brain tumor biopsy. **I)** Analysis of immune population through FAST cycling of biopsied cells correlates strongly with analysis with conventional flow cytometry.

### Biopsy does not lead to immune changes after two or five days

A previous *in vivo* imaging study showed that biopsy increased migration and proliferation of tumor cells locally in a glioma model, induced by infiltration of macrophages^16^. To assess the impact of biopsy on the immune response, we implanted tumors and biopsied mice as described above. Brains were harvested for flow cytometry analysis two or five days after biopsies or sham surgeries. No significant changes in leukocytes (See *Supplementary Table 1* for marker identification), macrophages, CCR2^+^ macrophages, monocytes, CCR2^+^ monocytes, microglia, T-cells, CD8^+^ T-cells, CD8^+^ PD-1^+^ T-cells, CD4^+^ T-cells, CD4/CD8 T-cells, and CD4^-^/CD8^-^ T cells were noted (*Figure 1C, D)*.

### Multiplexed cellular imaging accurately identifies cellular subsets in glioma

For multiplexed immune profiling in needle biopsy samples, we utilized ultra-fast cyclic immunolabeling method (FAST)^17^. FAST imaging exploits highly accelerated click reaction of bio-orthogonal tetrazine/trans-cyclooctene (Tz/TCO), which enables iteration of rapid quenching (>99% quenching in <10 seconds) and re-staining of immunolabeling while maintaining the integrity of single cells. A FAST probe is composed of a fluorophore (Alexa Fluor (AF)488, AF555, or AF647), a TCO for clicking with a Tz-labeled quencher (Black hole quencher, BHQ3), and a flexible polyethylene glycol -4 (PEG4) side chain for efficient antibody labeling (*Figure 1E*). The FAST probes were activated with N-Hydroxysuccinimide (NHS) esters and conjugated to antibodies against proteins of interest^17,18^. We designed a panel of FAST-antibodies against antigens most relevant in analyzing the microenvironment of GL261 tumors (*Supplementary Table 1*). A total of 22 markers were analyzed after 6 cycles of FAST imaging in each biopsy sample on a glass slide (*Figure 1F)*. Cell types were identified based on the fluorescent intensity of the protein markers in individual cells to determine the immune composition of each biopsy sample (*Figure 1G,H)*. After the biopsy, the tumors were extracted and processed for flow cytometry. The analysis result of FAST imaging of needle biopsies was then validated by flow cytometry analysis of the same tumor tissues *(Figure 1I*, R^2^=0.90), indicating that our FAST technique could accurately identify cellular population in this model.

### Serial biopsy shows changes in the immune profile over time

To test our set-up and evaluate the changes in the immune profile over time, we injected two groups of four mice each with GL261.Fluc.GFP cells, and confirmed tumor growth on day 10 with IVIS. Biopsies were taken on day 10, and day 12 or 15 after tumor inoculation (*Figure 2A)*. Between 5,000 and 25,000 cells were analyzed from each biopsy with multiplexed cellular imaging. We analyzed the changes in various innate and adaptive immune cell populations over two days (*Supplementary Figure 1A-T*), and over five days (*Supplementary Figure 2A-T*). Trends could be observed in the changes over time (*Figure 2B,C*). Differences in size and change in immune populations was observed among the different mice, further underscoring the notion that even in highly controlled mouse models, glioma cells and the immune response they induce are a heterogeneous and variable process. To visualize the changes over time, we pooled the data from day 0 and plotted the percentages of relevant populations observed at day two and five (*Figure 2D-O*). For statistical analysis, a paired t-test was performed on changes within the same mice (*Supplementary Figure 1 and 2*). From all biopsied cells, between 5% and 20% were CD45 positive, with a decreasing trend observed after 5 days (*Figure 2D*). The percentage of macrophages in these CD45 cells trended upward, while the percentage of monocytes trended downwards (*Figure 2E,F)*. The percentage of microglia, neutrophils, and natural killer (NK) cells was consistent over time (*Figure 2 G-I*). No change was observed in the percentage of dendritic cells (*Supplementary Figure 1&2 M)*. On average, the percentage of CD8 T cells in the tumor remained stable (*Figure 2J*), while the percentage of CD8 T cells expressing PD-1 (programmed cell death protein 1) increased significantly after 5 days (*Figure 2K, Supplementary Figure 2S).* No significant changes were observed in CD4 T cells and Treg cells, while an upward trend was observed in the percentage of B cells (*Figure 2L-N)*. The expression of PD-L1 on CD45 cells showed large variations among mice and time points (*Figure 2O*). Aside from PD-1 expression on CD8 T cells, other markers implicated in the immune response were also measured. Expression of CTLA-4 and IFN-! was assessed on CD8 T cells (*Supplementary Figure 1&2 Q-S),* CCR2 and IL-12 expression on macrophages *(Supplementary Figure 1&2 C,D)*, and IL-12 expression on microglia and dendritic cells (*Supplementary Figure 1&2 G, N).* IL-12 expression on dendritic cells appeared to diminish over time, but this change was not significant. Overall, gradual changes could be detected in the tumor (microenvironment) over time, most notable in PD-1 expression on CD8 T cells.

**Figure 2:**
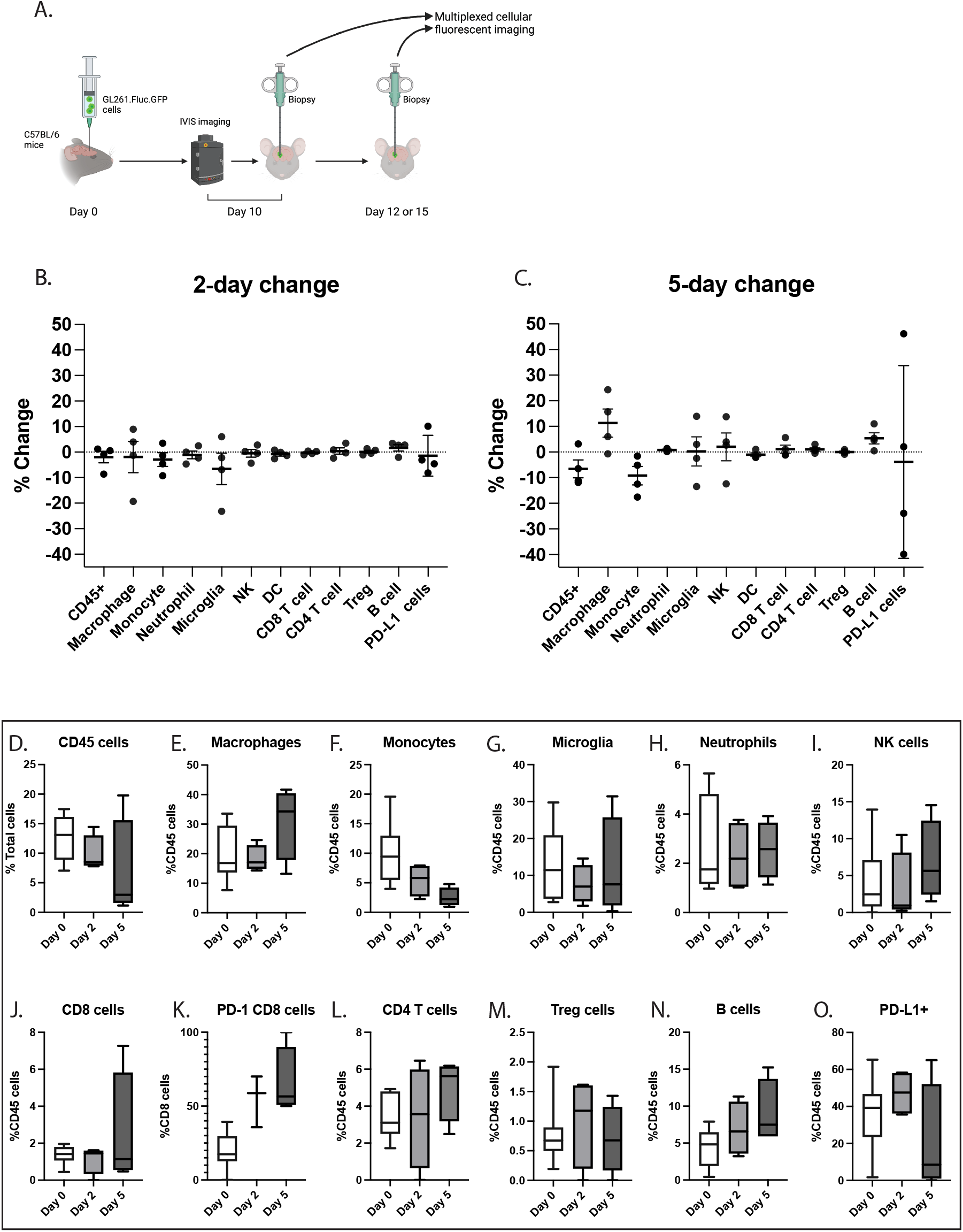
serial sampling of brain tumor shows changes in immune populations over time. **A)** Schematic flowchart of experimental set-up for repeat biopsy. Four mice per group with one group biopsied at day 10 and 12, and one group at day 10 and 15. **B)** Change in percentage of immune cells between biopsies per mouse after two days. Change in CD45 percentage as compared to total cells, change in other immune populations as compared to CD45 positive cells. **C)** Change in percentage of immune cells between biopsies per mouse after five days. Change in CD45 percentage as compared to total cells, change in other immune populations as compared to CD45 positive cells. **D-O)** Changes in immune populations between day 0 (n=8), day 2 (n=4, except PD-1 CD8 cells with n=3), and day 5 (n=4).

### Biopsy itself has a minor but significant effect on the local immune response

Biopsy and surgery can have an impact on the local immune response in glioma models^12,19^. To assess the impact of our biopsy model on the local immune response, we tested a new experimental set-up. Ten mice were injected with GL261.Fluc.GFP tumor cells and stratified into two groups of similar average tumor size as measured with IVIS on day 10 (*Figure 3A)*. Biopsies were performed at the same coordinates of tumor implantation in all mice. Five days after the initial biopsy, one group was biopsied again at the same coordinates of implantation while the other was biopsied at a different location in the tumor, 0.3mm away from the previous site in all three directions (X, Y, and Z), thus comparing the previously biopsied area with a non-biopsied area. A similar percentage of GFP positive tumor cells was observed in the same-site and different-site biopsies at day 5, indicating that the biopsies in both groups were taken from the tumor mass (20.5±5.9% in the same-site biopsies and average 22.2±6.3% in the different-site biopsies). We evaluated changes in immune populations in the tumor between the first and second biopsy in the samples taken from the same site (*Supplementary Figure 3A-T)* and those taken from different sites (*Supplementary Figure 4A-T*) to assess the impact of the biopsy itself over five days. We then compared the changes on immune populations between the two groups (*Figure 3 B,C*), highlighting the relevant changes in *Figure 3D-O.* The number of CD45^+^ cells was slightly higher on average in these samples compared to the previous experiments, between 20% and 50% (*Figure 3D,J),* but did not differ between the same site and different site. The percentage of CD45 cells expressing PD-L1 was significantly lower in non-biopsied area in the tumor (*Figure 3E*, p=0.04), while remaining the same in a previously biopsied area (p=0.7) (*Figure 3K)*. The presence of neutrophils increased significantly at the site in which a previous biopsy had occurred (p=0.04), while remaining stable at a different site in the tumor (p=0.4) (*Figure 3F,L*). In both groups the number of CD4 T cells seemed to decrease over time, although it was only significant in the different site biopsy group (p=0.046 vs. p=0.84) (*Figure 3G,M*). As in the previous experiment, the number of T cells expressing PD-1 increased over time in both groups, but the increase was not significant in the group biopsied at a different site (p= 0.012 versus p=0.054, respectively) (*Figure 3H,N*). Finally, a significant decrease in NK cells was observed in the same-site biopsy group, but not in the different-site group (p=0.02 versus p=0.3) (*Figure 3 I,O*). Overall, minor changes were observed between the previously biopsied and surgery-naïve site, with biopsy driving expression of PD-L1, presence of neutrophils and PD-1 expression on CD8 T cells.

**Figure 3:**
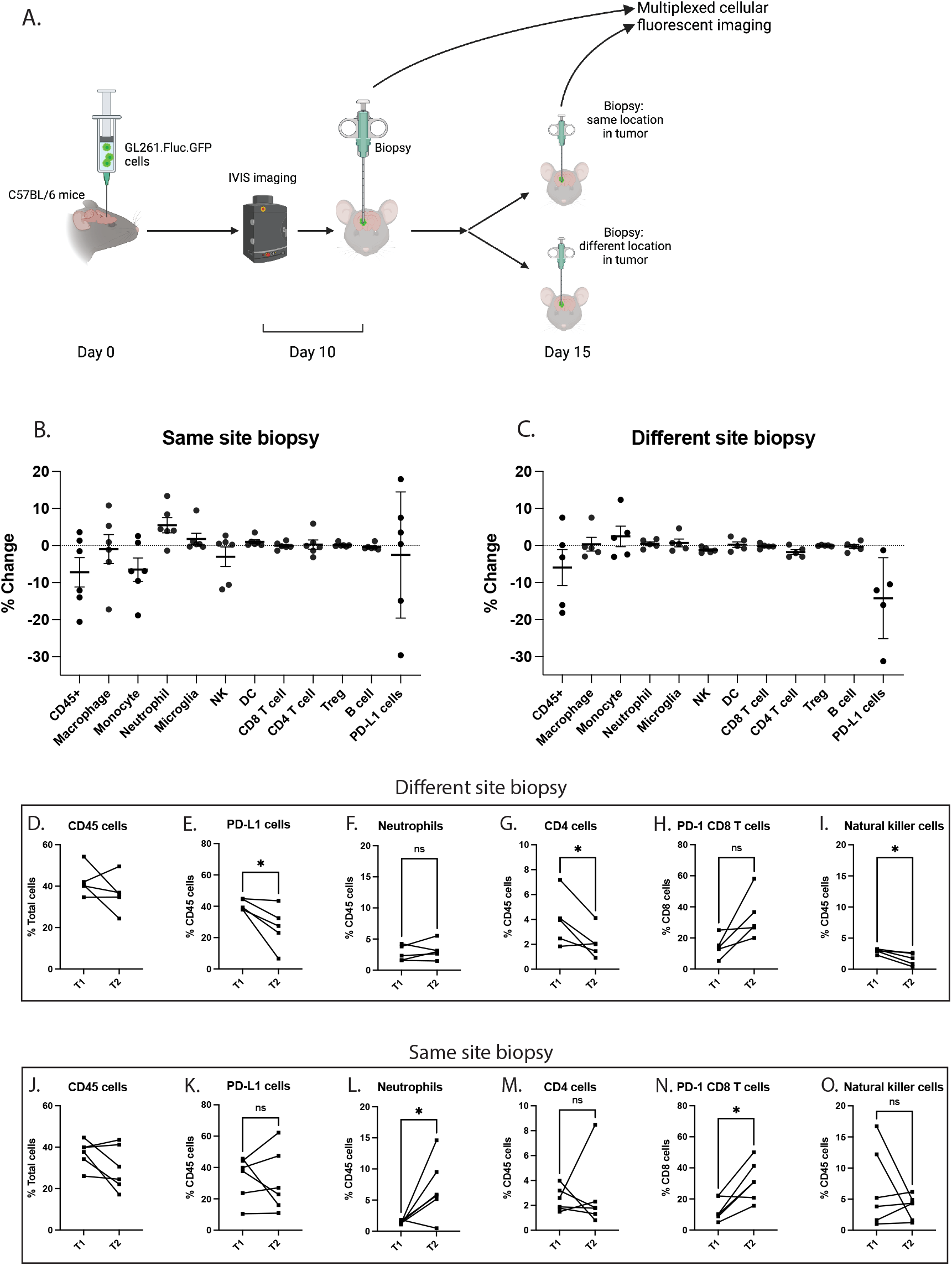
repeat sampling of tumor site induces minor changes. **A)** Schematic flowchart of experimental set-up for repeat biopsy. Six mice per group with one group biopsied at the exact same site twice, and one group biopsied in a different location. **B)** Change in percentage of immune cells per mouse in same-site biopsy. Change in CD45 percentage as compared to total cells, change in other immune populations as compared to CD45 positive cells. **C)** Change in percentage of immune cells per mouse in different-site biopsy. Change in CD45 percentage as compared to total cells, change in other immune populations as compared to CD45 positive cells. **D-I)** Changes in immune populations per mouse between day 0 (T1) and day 5 (T2) in different-site biopsy. Student’s t-test. * p<0.05. ns: not significant. **J-O)** Changes in immune populations per mouse between day 0 (T1) and day 5 (T2) in same-site biopsy. Student’s t-test. * p<0.05. ns: not significant

## Discussion

In this study, we demonstrate the feasibility of repeat biopsy in a mouse model of glioma, and illustrate the changes in the immune response over time. Survival of mice was not impacted due to biopsy. Immune profiling through ultra-fast cycling with multiplexed cellular imaging accurately identified the immune population present and showed changes in the immune response over time. Biopsy did not alter the overall immune response in the tumor, but locally some changes could be observed after five days.

Different forms of biopsy exist to allow for the safe retrieval of cells. Image, guided or stereotactic needle placement can be used to extract single cells, cell clusters (occasionally termed fine needle aspirate or FNA), microcores or conventional cores of tissue. Larger biopsies can provide information on heterogeneity and spatial biology, but carry more risk, especially when retrieving cells from vital structures such as the brain^5^. Repeated biopsies are therefore preferably performed by low-impact forms of biopsy such as FNA. To allow for thorough analysis of only a few numbers of cells, multiplexed cellular imaging with FAST staining has been developed^17,18^. FAST staining is compatible with all forms of biopsies, both of histologic tissue and single cells. Here we demonstrate its potential in monitoring immunological changes in the brain and specifically in glioma over time, showing a strong correlation to immunoprofiling via flow cytometry. Compared to other studies of GL261 in mice, we identified similar percentages of immune populations in the CD45 compartment, although the number of T cells appeared to be lower^11,20^. Most T cells are found on the tumor border or in the perivascular region^21^, and are included when the whole tumor is analyzed, while our biopsies are taken in the tumor core where T cell infiltration is possibly less abundant^22^. There are a number of benefits to studying the changes in the immune system via our method. It removes the possible confounding differences between mice and batches of cells, as changes over time and in response to treatment can be studied within the same mouse. Even though mouse glioma models have been standardized, heterogeneity due to mutations in the cell lines, handling of animals, cell culture passaging, and various other external factors tend to cause differences in tumor progression and the immune response among experiments^23,24^. Repeat biopsy can correct for that by evaluating changes in the same mouse and tumor. The number of animals needed to study these changes can thus also be reduced. Multiplexed cellular imaging decreases the numbers of cells needed for extensive immune profiling, allowing for small biopsies. While in the present study we focused exclusively on immune profiling through multiplexed cellular imaging, studying changes in other cellular profiles such as gene expression, protein expression or metabolomics, can also be studied via this biopsy method.

The primary goal of this study was to design and evaluate serial biopsy and multiplexed cellular imaging with FAST in a glioma model in mice. While the preliminary data of the changes in immune populations shows interesting results, we are cognizant of the limitations in interpreting this data. Briefly, we observed an increase in PD-1 expression on CD8 T cells over time, enhanced through biopsy, while the number of CD8 T cells in the tumor did not increase. PD-1 is a protein expressed on T cells during T cell activation, and has a role in suppressing the T cell response^25^. In patient and mouse samples of glioma, PD-1 is expressed on 50% to 90% of tumor infiltrating CD8 T cells^13,26,27^, comparable to our data at the last biopsy point. PD-1 expression in glioblastoma has been related to T cell exhaustion, as it is correlated with high expression of other markers for exhaustion, such as TIM-3 and Lag-3^26–28^. Expression of CTLA-4, another marker for T cell exhaustion^29^, did not increase over time and was not as strong compared to PD-1 in our data, in line with other studies^11,26,27^. Otvos et al. previously showed that surgical resection of CT2A in mice induces upregulation of PD-1 expression on CD8 T cells at the tumor border and in the bone marrow^12^. While exposure to the tumor seems to be the biggest driver for PD-1 expression, tissue injury through surgery or biopsy may further drive PD-1 expression and T cell exhaustion. Our results also showed changes in CD4 expression, NK-cells and PD-L1 over time and in response to biopsy, but more studies need to be done to confirm the validity of these results.

A topic of future investigations is whether changes between biopsied and non-biopsied areas of the tumor are sustained over time, and how other factors influence these changes. Dexamethasone, a corticosteroid often used in glioblastoma patients, has been shown to repress biopsy-induced changes^16^. This study indicates that performing biopsies at different locations in the tumor may be preferable, to minimize confounding variables from biopsy induced changes.

We are cognizant of several limitations to our method. Aiming for different areas in the tumor introduces spatial heterogeneity in the sampling and increases the risk of missing the tumor during biopsy. However, our stereotactic frame allowed for precise measurement and positioning of the biopsy needle, and all our samples contained similar percentages of tumor cells. Another factor we cannot control for is contamination from immune cells in the vasculature. As the biopsies are done in live animals, the mice are not perfused and contamination from the blood stream is possible. No hemorrhages in the brain were observed however during biopsy and upon necropsy, and similar immune populations were found between whole tumor, sodium-chloride perfused samples, and biopsy acquired, non-perfused, samples.

Here we demonstrate the possibilities of repeat biopsies in mice combined with multiplexed cellular imaging. To provide clearer insight into changes over time and the response to the biopsy itself, multiple different glioma models should be evaluated over longer periods of time. While this techniques works well in mice, translation to the clinic is difficult as biopsy is associated with increased mortality and morbidity^5^. In superficial tumors, such as melanomas or squamous head-neck tumors^18^ repeat biopsy may be more practical. Multiplexed cellular imaging can be utilized in instances when little material is available for analysis.

Overall, we demonstrate the feasibility of our technique in mice, and demonstrate subtle changes in the tumor microenvironment which could indicate increased T cell exhaustion over time.

## Methods and Materials

### Cell culture

Mouse glioma cell line GL261 (NCI GL261 0507814) was stably transduced with the CSCW2.Fluc.IRES.GFP lentivirus, kindly provided by Dr. Miguel Sena-Esteves (UMass Medical Centre).

### Mice

C57BL/6 mice (Charles River Laboratories Wilmington, MA), aged 8-12 weeks both males and females were used for experiments. Animals were housed at the animal facility of the Massachusetts General Hospital under standard laboratory conditions, with free access to water and food. After tumor cell injection, mice were monitored daily to assess health, appearance and behavior. All experiments were approved under IACUC protocol 2009N000054.

### Tumor cell implantation

Mice were anesthetized with isoflurane, mixed with oxygen at 3.5% for induction and 2% for maintenance. Subcutaneous buprenorphine was given for analgesia. A burrhole was made on the left side, 2.0mm anterior and 0.5mm lateral of Bregma. 100,000 cells suspended in 2μl OptiMEM (Thermo-Fisher) were implanted in the left striatum at a depth of 2.5mm from the skull surface. After injection, the incision was closed with 5/0 vicryl sutures.

### Bioluminescence

100 μl of D-Luciferin (Thermo-Fischer) (25 mg/ml in saline) was injected intra-peritoneally in isoflurane anesthetized mice. After 10 minutes the mice were imaged using the In Vivo Imaging System (IVIS) Spectrum connected to XGI-8 Anaesthesia System (PerkinElmer, Waltham, MA, USA). Bioluminescence was expressed as Total Flux per second.

### Biopsy procedure

Mice were anesthetized with isoflurane and buprenorphine, as described above. The surgical site was cleaned and the old incision re-opened. For the biopsy, a 22s gauge Hamilton syringe with rubber plunger was used. Three biopsies of around 5 ul were taken from the tumor core by aspirating 10 ul cells at the coordinates of the original injection site. Hemostasis was achieved with saline irrigation and sterile cotton swabs, and the incision was closed with 5/0 vicryl sutures. After 2 or 5 days, the procedure was repeated as described above. For the control group, the procedure was performed as described, with omission of the actual biopsy. Anesthesia time was matched to those undergoing biopsy. For the experiment evaluating the effects of the biopsy itself, biopsies were performed on all mice on day ten at the site of tumor implantation. As measured from bregma: 2.0 mm anterior, 0.5 mm lateral, and 2.5 mm deep. On day five, biopsies were repeated at either the exact same coordinates from bregma, or a new burr hole was drilled at 1.7 mm anterior, 0.8 mm lateral with biopsies taken at 2.2 mm deep to avoid the preciously biopsied area.

### Brain harvesting and single-cell preparation

Two or five days after biopsy, the mice were deeply anesthetized with a mixture of ketamine (17.5 mg/ml) and xylazine (2.5 mg/ml) followed by trans-cardial perfusion with 50 mL Dulbecco’s phosphate buffered saline without Mg^2+^ and Ca^2+^ (DPBS) using a perfusion pump (Minipump Variable Flow, Fisher Scientific). The brains were harvested and the tumors excised with a small margin of normal white matter. The tissue was then processed into single-cell suspensions for cell sorting and mass cytometry analysis using the Tumor Dissociation Kit (130-096-730) from Miltenyi (Bergisch Gladbach, Germany) according to manufacturer’s instructions. Tumor were cut into small pieces and further dissociated with the gentleMACS™

Dissociator (Miltenyi) and enzymes. Myelin was removed by resuspending the cells in Myelin Removal Beads II (130-096-733, Miltenyi) and running them over magnetic columns. Finally, red blood cell lysis was performed with mouse RBC lysis buffer (Boston Bioproducts, Ashland, MA, USA).

### Staining for whole tumor flow cytometry

Cells were blocked with TruStain fcX™ (anti-mouse CD16/32, BioLegend, #101319, clone 93, 1:100) for 10 minutes on ice. Cells were then stained for 10 minutes on ice. The following antibodies were used: anti-CD45-BV605 (BioLegend, 103139, clone 30-F11, 1:100), anti-CD11b-PE-Cy7 (Biolegend, 101215, clone M1/70, 1:100), anti-CD3-PerCP-eFluor 710 (Thermo Fischer, 46-0032-82, 1:50), anti-CD8-BV785 (Biolegend, 46-0032-82, clone 53-6.7, 1:50), anti-CD4-alexa fluor 700 (Biolegend, 100430, clone GK1.5, 1:50), anti-Ly6C-PE (Biolegend, 1280008, clone HK1.4, 1:200), anti-CCR2-APC (Biolegend, 150628, clone SA203G11, 1:25). Live-dead staining was done with Zombie UV (Biolegend, 423107) After staining, cells were centrifuged at 400 x G for 8 min, resuspended in PBS with 0.5% BSA and passed through a 35 μm nylon mesh strainer (BD Falcon). Flow analysis was done on the BD Fortessa X-20).

For flow cytometry analysis for validation of immune profiling of needle biopsies by FAST imaging, the same tumors from which the biopsies were obtained were extracted. Samples were first incubated with True Stain fcX^TM^ antibody (anti-mouse CD16/32, BioLegend, #101319, clone 93, 1:100) in PBS containing 0.5% BSA and 2 mM EDTA before staining with antibodies directly conjugated to fluorophores for flow cytometry. Propidium iodide was used to exclude dead cells. In addition to FAST-labeled antibodies listed in Supplementary Table 2, the following antibodies were used: anti-CD3e (clone 145-2C11, Biolegend), CD4 (clone GK1.5, BD Biosciences), CD45 (clone 30-F11, Biolegend), CD8a (clone 53-6.7, Biolegend), CD11b (clone M1/70, Biolegend), CD11c (clone N418, Biolegend), F4/80 (clone BM8, Biolegend), MHC II (clone M5/114.15.2, Biolegend), CD25 (clone PC61, Biolegend), FoxP3 (clone MF-14, Biolegend), Ly6C (clone HK1.4, Biolegend), Ly6G (clone 1A8, Biolegend), and TCRb (clone H57-597, Biolegend) were used for validation of marker staining.

### Synthesis of fluorochrome/quencher pair

FAST probes are built around a modular linker between fluorochromes and antibodies with an embedded TCO for clicking with a tetrazine-quencher. FAST probes were custom synthesized at large scale as described in detail in our previous study, stored as the carboxylic acids, and activated for antibody labeling with our in-situ activation chemistry, which eliminates the need for purification and long-term storage of the NHS esters^17,18^. The dTCO-PEG6-CO2H blocking reagent was synthesized from dTCO-PNP and amino-dPEG6-CO2H and characterized by LCMS, as described in *Figure 1E*. All reagents were obtained from commercial sources at the highest grade available and used without further purification. Fluorophores were purchased from Click Chemistry Tools or Fluoroprobes. BHQ®-3 Amine was purchased from LGC Biosearch Technologies. N-⍺-Boc-N-ε-Fmoc-Lysine was purchased from Chem-Impex. Amino-dPEG®n-carboxylic acids (n=4,6) were obtained from Quanta BioDesign. Dry solvents and coupling reagents were obtained from Sigma Aldrich. rTCO-PNP and dTCO-PNP were a generous gift of Dr. Hannes Mikula (TU Wien, Austria).

### FAST antibody conjugation

Antibodies without carrier were purchased (*Supplementary Table 1*) to be labeled with FAST probes as previously described^18^. A Zeba column (Thermo Fisher) was used to buffer-exchange the antibodies into bicarbonate buffer (pH 8.4). Then antibodies (1-3 mg/ml) were incubated with a 5- to 10-fold molar excess of the FAST probe with 10% DMSO for 25 minutes at room temperature in dark. For desalting and removal of unreacted dye molecules after the conjugation reaction, FAST-antibody conjugate was loaded onto another 40k Zeba column equilibrated with PBS. The absorbance spectrum of the FAST-labeled antibody was measured using a Nanodrop 1000 (Thermo Scientific) to determine the degree of labeling (DOL). The known extinction coefficients of the dye (AF488, AF555, AF647), IgG antibody, and correction factor for the dye absorbance at 280nm were used for calculation of DOL. The FAST-labeled antibodies were stored protected from light at 4°C in PBS. Information of the antibodies used for the single-cell profiling of clinical samples are summarized in *Supplementary Table 1*. Antibodies were tested and validated on positive cell lines, mouse splenocytes or mouse brain tissue before usage.

### FAST imaging

Cells were fixed for 10 minutes in 4% PFA and permeabilized for 25 minutes with 0.5% Triton-X100 prior to staining. Immunostaining for FAST imaging was performed in accordance with typical immunofluorescence protocols. After blocking with Intercept Blocking buffer (LI-COR Biosciences) for 30 minutes, cells were stained with FAST-conjugated antibodies. Antibodies were diluted to 2-5 µg/ml in Intercept Blocking buffer before staining. Stained cells were washed 3 times with PBS before imaging. Following image acquisition, cells were briefly incubated with 10 µM Tz-BHQ (<10 seconds) in PBS-bicarbonate buffer (pH 9) for quenching. Free Tz-BHQ was removed by 3 washes with PBS-bicarbonate buffer, and the cells were imaged again in the same fields of view to record the quenched signal. Before antibody staining of the subsequent cycle, cells were briefly incubated in a solution of 20 µM dTCO-PEG6-CO2H in order to block any residual Tz-BHQ3 from reacting with FAST antibodies of the next cycle. The same staining, imaging, and quenching cycle was repeated until all of the target proteins were imaged. A fraction of the sample was set aside and incubated with isotype antibodies as a negative control. The controls were imaged every cycle following the same protocol as described above.

### Fluorescence microscopy

An Olympus BX-63 upright automated epifluorescence microscope was used to acquire fluorescent images. DAPI, FITC, Cy3, and Cy5 filter cubes were used to image DAPI nuclear stains, AF488, AF555, and AF647 fluorophores, respectively. Depending on the density of cells on the glass slides, 15-30 fields of view were imaged for each specimen to capture a total cell population sufficient for analysis. X-Y coordinates for each field of view were saved to enable automatic imaging of the same set of cells in every cycle using Multi Dimensional Acquisition in Metamorph software.

### Imaging analysis

CellProfiler^30^ was used for image registration, cell segmentation, and measurement of the fluorescent intensities in individual cells. Acquired images were corrected for an illumination function and aligned using normalized cross correlation to compensate for minor pixel shifts that occur during cyclic imaging. For background correction, fluorescence signals measured in quenched images were subtracted pixel-by-pixel from the immuno-stained images of the following cycle. Cells were identified using the DAPI signal of the last image cycle as input, and the cell boundaries were segmented using maximum projection of intensity measured in all channels. A small fraction of cells (typically well below <10%) that were lost during repeat imaging cycles were excluded from the image analysis. In every identified cell, an areal mean fluorescence intensity was used for subsequent computational analyses.

### Statistics

GraphPad Prism was used for statistical analysis. Results were expressed as mean ± SEM. Statistical tests included one-way ANOVA followed by Tukey’s or Dunnett’s multiple comparison test. When applicable, the unpaired one-tailed and two-tailed Student’s t tests using Welch’s correction for unequal variances were used.

## Funding

National Institute of Health grant R35 CA232103 (TSvS, EA, XOB)

National Institute of Health grant R01CA257623 (JO, PK, RW)

National Institute of Health grant P01CA240239 (JO, PK, RW)

CSB Development Grant (JO, RW)

Bontius Stichting (TSvS)

the Nijbakker-Morra Fund(TSvS)

Foundation Vrijvrouwe van Renswoude (TSvS)

Bekker-la Bastide Fund (TSvS)

## Author Contributions

Conceptualization: MLDB, RW, XOB

Methodology: TSvS, JO, EA, PK

Investigation: TSvS, JO

Visualization: TSvS, JO

Supervision: XOB, RW, MLDB

Writing—original draft: TSvS, JO

Writing—review & editing: EA, PK, XOB, RW, MLDB

## Data and materials availability

The complete dataset of immunoprofiling data is available upon request to the corresponding authors.

