## Supplemental materials for "Probing the glioma micro-environment: analysis using biopsy in combination with ultra-fast cyclic immunolabeling"

#### Supplementary Figure 1: two day biopsy

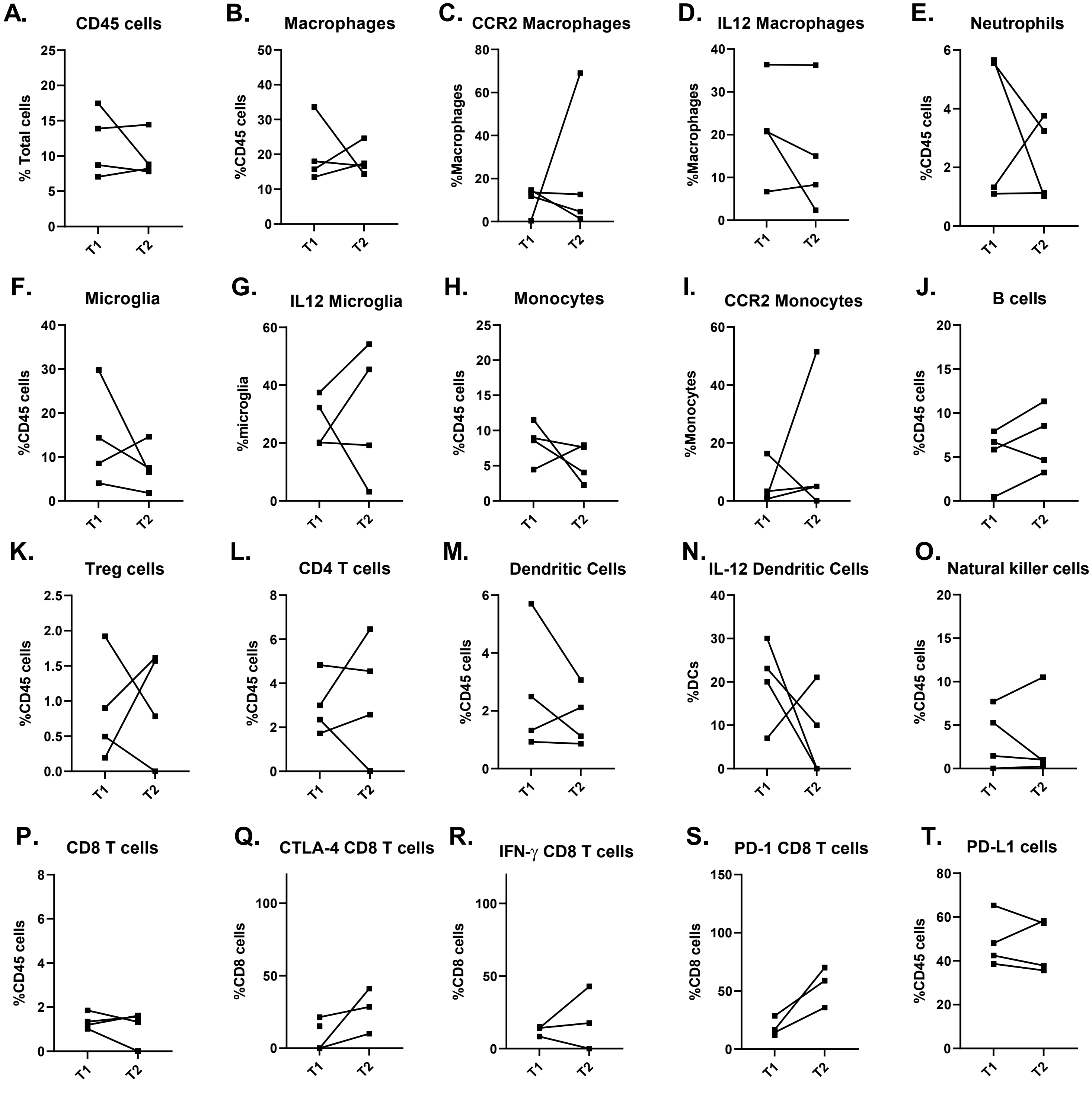

#### Supplementary Figure 2: five day biopsy

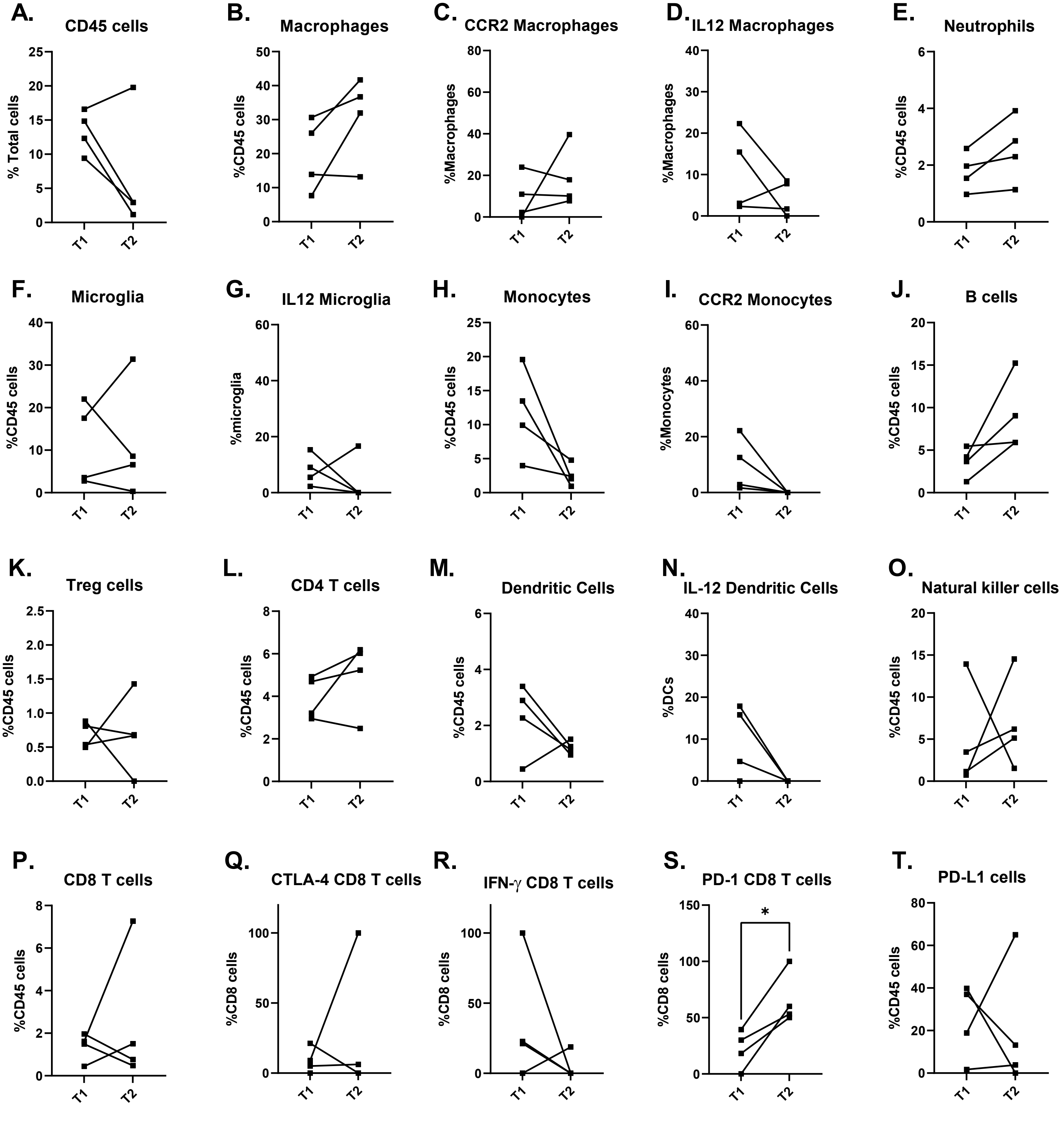

#### Supplementary Figure 3: 5-day biopsy, same site

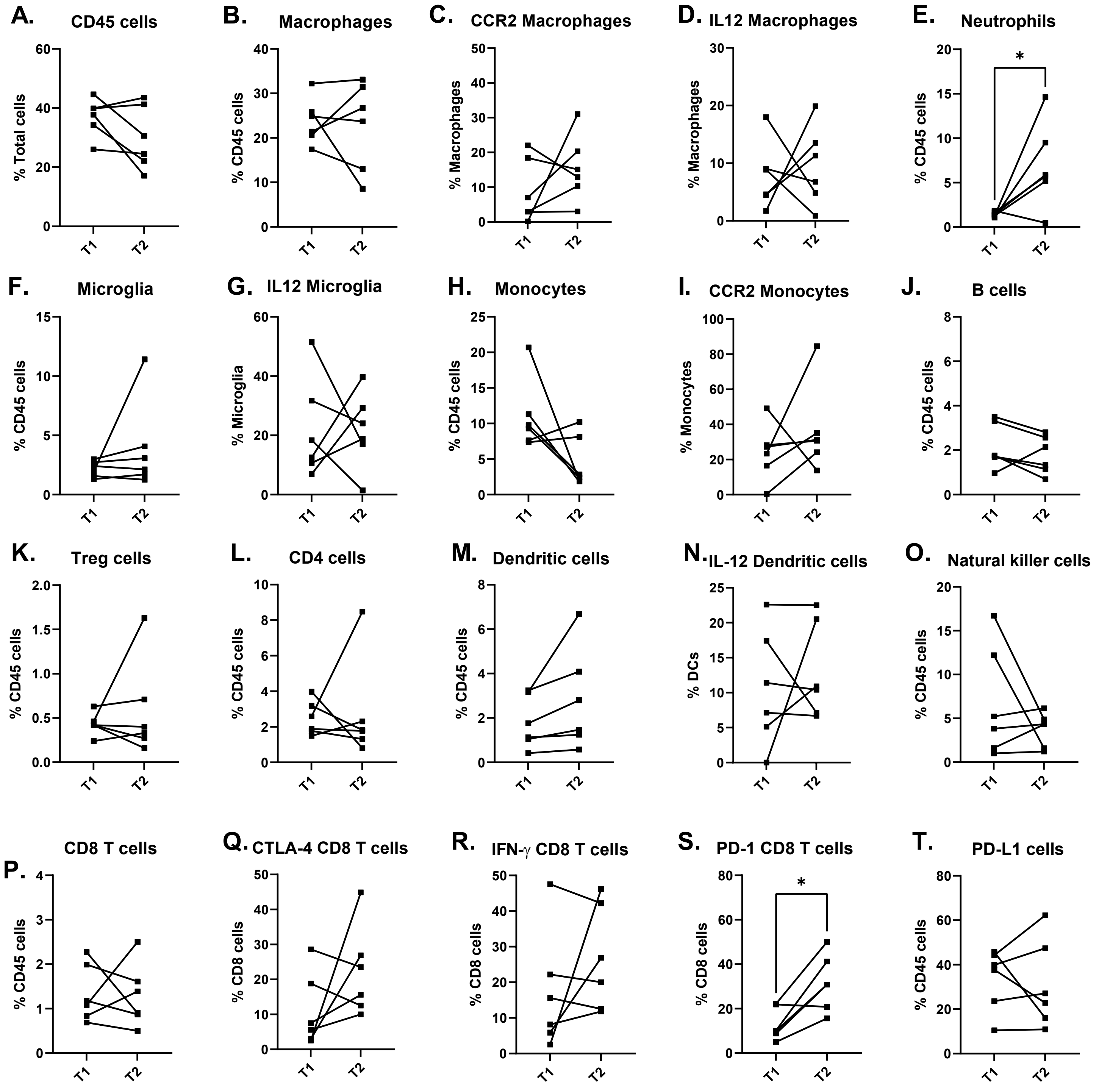

### Supplementary Figure 4: 5-day biopsy, different site

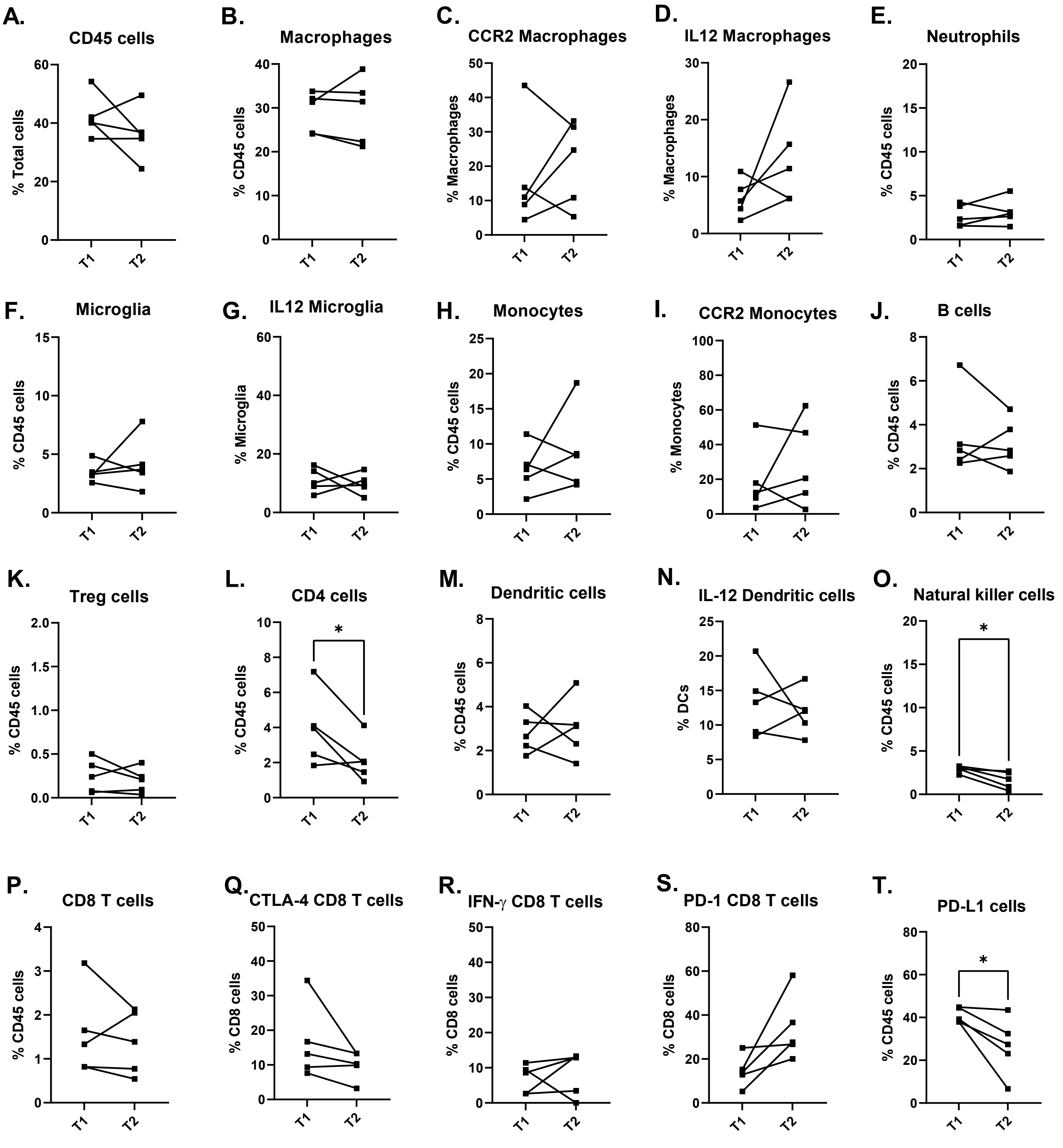

| **Supplementary Table 1:** Markers for cell subtyping in *Figure 1 C,D* after excluding doublets | |
|---|---|
| Leukocytes | $CD45^{high}$ |
| Macrophages | $CD45^{high}$/$CD11b^{high}$/$Ly6C^{low}$ |
| $CCR2^{+}$ Macrophages | $CD45^{high}$/$CD11b^{high}$/$Ly6C^{low}$/$CCR2^{high}$ |
| Monocytes | $CD45^{high}$/$CD11b^{high}$/$Ly6C^{high}$ |
| $CCR2^{+}$ Monocytes | $CD45^{high}$/$CD11b^{high}$/$Ly6C^{high}$/$CCR2^{high}$ |
| Microglia | $CD45^{dim}$ |
| T-cells | $CD45^{high}$/$CD3^{high}$ |
| $CD8^{+}$ T-cells | $CD45^{high}$/$CD3^{high}$/$CD8^{high}$/$CD4^{low}$ |
| $CD8^{+}$ $PD1^{+}$ T-cells | $CD45^{high}$/$CD3^{high}$/$CD8^{high}$/$CD4^{low}$/$PD1^{high}$) |
| $CD4^{+}$ T-cells | $CD45^{high}$/$CD3^{high}$/$CD4^{high}$/$CD8^{low}$ |
| CD4/CD8 T-cells | $CD45^{high}$/$CD3^{high}$/$CD4^{high}$/$CD8^{high}$ |
| $CD4^{-}$/$CD8^{-}$ T cells | $CD45^{high}$/$CD3^{high}$/$CD4^{low}$/$CD8^{low}$ |

**Supplementary Table 2: Antibodies for FAST imaging of mouse immune cell populations.**

| Markers | Target population | Clone | Source | Catalog # | Dye |
|---|---|---|---|---|---|
| CD45 | Hematopoietic cells | 104.2 | Bio X Cell | BE0300 | AF488 |
| CD3 | T cells | 145-2C11 | Biolegend | 100306 | AF647 |
| CD8 | CD8+ T cells | 53-6.7 | Bio X Cell | BE0004-1 | AF555 |
| CD4 | CD4+ T cells | GK1.5 | Bio X Cell | BE0003-1 | AF647 |
| FoxP3 | Regulatory T cells | MF-14 | Biolegend | 126402 | AF647 |
| NK1.1 | NK cells | PK136 | BD Pharmingen | 553162 | AF555 |
| CD19 | B cells | 1D3 | BD Pharmingen | 553783 | AF488 |
| CD20 | B cells | SA271G2 | Biolegend | 152102 | AF488 |
| CD11b | Myeloid cells | M1/70 | Bio X Cell | BE0007 | AF647 |
| F4/80 | Macrophages | CI:A3-1 | Bio X Cell | BE0206 | AF555 |
| CD11c | Dendritic cells | N418 | Biolegend | 117302 | AF647 |
| Ly6G | Neutrophils | 1A8 | Bio X Cell | BE0075-1 | AF647 |
| Ly6C | Monocytes | Monts1 | Bio X Cell | BE0203 | AF488 |
| MHCII | Dendritic cells | M5/114 | Bio X Cell | BE0108 | AF488 |
| PD-1 | Various | 29F.1A12 | Bio X Cell | BE0273 | AF555 |
| PD-L1 | Various | 10F.9G2 | Bio X Cell | BE0101 | AF555 |
| CTLA4 | Various | UC10-4F10-11 | Bio X Cell | BE0032 | AF488 |
| IFN-ɣ | Cytotoxic T cells | XMG1.2 | Bio X Cell | BE0055 | AF555 |
| IL-12β | Dendritic cells | C17.8 | Bio X Cell | BE0051 | AF488 |
| CCR2 | Macrophages/Microglia | 475303 | R&D Systems | MAB55382 | AF647 |
| TMEM119 | Microglia | 28-3 | Abcam | Ab209064 | AF647 |
| P2RY12 | Microglia | 4H5L19 | Invitrogen | 702516 | AF555 |
| Anti-Rabbit-IgG | Secondary antibody | 6B9G9 | Biolegend | 410404 | AF647 |
